# Complementing the EGFR dynamic interactome using live-cell proximity labeling

**DOI:** 10.1101/2022.02.02.478771

**Authors:** Charlotte A.G.H. van Gelder, Wouter van Bergen, Pieter C. van Breugel, Maarten Altelaar

**Affiliations:** Biomolecular Mass Spectrometry and Proteomics, Bijvoet Center for Biomolecular Research and Utrecht Institute for Pharmaceutical Sciences, Utrecht University, Padualaan 8, 3584 CH Utrecht, The Netherlands; Netherlands Proteomics Center, Padualaan 8, 3584 CH Utrecht, The Netherlands

**Keywords:** Live cell proximity labeling, APEX2 - EGFR, Proteomics, Differential trafficking

## Abstract

The epidermal growth factor receptor (EGFR) is a member of the receptor tyrosine kinase family (RTK) of transmembrane receptors, known to regulate many key cellular processes, including growth, proliferation, and differentiation. Its expression, activation, trafficking, and degradation have been extensively studied, as dysregulation of EGFR activation has been linked to a vast number of cancers. Activation of EGFR by different ligands results in distinct cellular responses, and the relative distribution of EGFR in different endosome pools in a process called endosomal sorting, leading to lysosomal degradation, or cell surface recycling, respectively, is considered a fundamental process in EGFR stimulation outcome. The EGFR interactome is therefore an essential element in the study of RTK functional selectivity. Here, we aimed to complement the existing EGFR interactome with spatio-temporal information on EGFR, its interactors, and phosphorylation state. We identified and quantified EGFR stable and transient interactions at different time points after stimulation using an EGFR-APEX2 fusion construct expressed in HEK293T cells and were able to use bystander proteins to map EGFR subcellular location at each time point. Utilizing the fast and concise biotinylation of proximity proteins by APEX2, we were able to detect slight differences in early signaling kinetics between TGF-α and EGF, thereby increasing our knowledge on RTK signaling and differential trafficking.

## Introduction

The epidermal growth factor receptor (EGFR) is a member of the receptor tyrosine kinase (RTK) family of transmembrane receptors, known to regulate many key cellular processes, including growth, proliferation, and differentiation ^1^. Its expression, activation, trafficking, and degradation have been extensively studied, as dysregulation of EGFR activation has been linked to a vast number of cancers ^1,2^. Because of this, EGFR has become the model receptor, also representing lesser-studied growth factor RTKs ^1^.

On the extracellular domain, seven ligands are known to activate EGFR, resulting in receptor dimerization, autophosphorylation of the intracellular kinase domains, and subsequent internalization ^1,3^. Interestingly, activation of EGFR by different ligands results in distinct cellular responses, which are not only regulated via differences in signal duration ^4,5^, but also modulation of protein-protein interactions (PPIs) ^6^, and the subcellular localization of the activated receptor ^7–9^. More specifically, the relative distribution of EGFR in different endosome pools in a process called endosomal sorting, leading to lysosomal degradation, or cell surface recycling, respectively, is considered a fundamental process in EGFR stimulation outcome ^3^.

The EGFR interactome is therefore an essential element in the study of RTK functional selectivity. Classical affinity purification mass spectrometry (AP-MS) approaches, mainly antibody-based, are limited in detecting transient interactions, and do not convey any information on spatial and temporal behavior of the protein of interest, as they are often performed under non-physiological conditions ^10,11^. Biotin-based proximity-labeling approaches, where the protein of interest is fused to either a promiscuous biotin ligase or an engineered ascorbate peroxidase, can overcome these challenges ^10,11^. In APEX2, an engineered ascorbate peroxidase is fused to the protein of interest and introduced into the model system. Activation of APEX2 results in the rapid formation of biotin-phenoxy radicals within a selective 20 nm labeling radius. This fast and precise labeling in living cells allows for the generation of time-resolved ‘snapshots’ of the protein-of-interest’s transient protein interactions, as well as subcellular location via the use of cellular compartment-specific proteins, so-called ‘bystander proteins’ ^12–14^.

Numerous studies have investigated the EGFR interactome upon stimulation ^15,16^, including Francavilla *et al, who* published a time-resolved analysis of EGFR signaling using a multilayered proteomics approach to study interactome, phosphoproteome, ubiquitinome, and late proteome in response to differential activation with tumor growth factor alpha (TGF-α), and epidermal growth factor (EGF), respectively ^3^. Moreover, a recent study by Ke *et al* investigated the spatiotemporal behavior of the EGFR signaling component STS1 ^17^. Here, we aimed to complement the existing EGFR interactome with spatio-temporal information on EGFR, its interactors, and phosphorylation state. We identified and quantified EGFR stable and transient interactions at different time points after stimulation using an EGFR-APEX2 fusion construct expressed in HEK293T cells and were able to use bystander proteins to map EGFR subcellular location at each time point. Utilizing the fast and concise biotinylation of proximity proteins by APEX2, we were able to detect slight differences in early signaling kinetics between TGF-α and EGF, thereby increasing our knowledge on RTK signaling and differential trafficking.

## Materials and Methods

### Cell culture

Human embryonic kidney 293 T-antigen (HEK293T) cells (ATCC) were cultured in Dulbecco’s modified Eagle’s medium (DMEM, Lonza) and supplemented with 10% fetal bovine serum (Thermo Fischer Scientific), 1% penicillin/streptomycin and L-glutamine in a humidified atmosphere with 5% CO_2_ at 37 °C.

### cDNA constructs and transient transfections

EGFR-GFP was a gift from Alexander Sorkin ^18^ (Addgene plasmid #32751). To generate the EGFR-APEX2 plasmid, the GFP moiety of pEGFR-eGFP was digested out with AgeI and NotI. APEX2, flanked with MreI and NotI restriction sites, was amplified using pcDNA3 Connexin43-GFP-APEX2 (Addgene #49385) as template. The resulting PCR product was digested with MreI and NotI and ligated into AgeI and NotI digested EGFR-eGFP, generating EGFR-APEX2. cDNA was introduced in HEK293T cells using jetPRIME transfection reagent (Polyplus) following the manufacturer’s protocol. In short, cells were plated at a confluency of 50-60%. For a 6-well plate format, 1 μg of DNA was introduced in 200 μl of jetPRIME buffer and 2 μl of jetPRIME reagent. Experiments were performed 24 to 48 hours after transfection.

### Medium starvation and EGFR stimulation

Cells were starved 30 minutes before stimulus with EGF (Merck) (100ng/mL) or Human TGF-α (Peprotech) (100ng/mL) at 37°C. Starvation medium consists of DMEM supplemented with 1% penicillin/streptomycin and L-glutamine. If stimulation with a ligand was followed by a labeling reaction, the starvation medium was also supplemented with 500 μM biotin-phenol (Iris Biotech).

### APEX reaction and cell lysis

Cells were incubated with biotin phenol (BP, Iris Biotech) supplemented DMEM at a final concentration of 500 μM for 30 minutes at 37°C/5%CO_2_. The APEX reaction was performed by introduction of 100 mM H_2_O_2_ (Merck) diluted in Dulbecco’s phosphate buffered saline (DPBS, Lonza) to a final concentration of 1 mM for 60 seconds at room temperature. The reaction was quenched by addition of ice-cold quencher solution, consisting of 10 mM sodium ascorbate (Sigma Aldrich), 5 mM Trolox (Sigma Aldrich), and 10 mM sodium azide (Sigma) in DPBS for 20 minutes on ice. Cell pellets were collected and lysed in RIPA lysis buffer (50 mM TRIS-HCl pH 7.4, 150 mM NaCl, 0.1% sodium dodecyl sulfate, 0.1% sodium deoxycholate, and 1% Triton X-100) supplemented with 10 mM sodium ascorbate, 5 mM Trolox, 10 mM sodium azide, 1 mM PMSF (Sigma), and complete mini EDTA-free protease inhibitor cocktail (Roche). Cell lysates were sonicated for 12 rounds of 5s (Bioruptor, Diagenode) and spun down at 14,000 rpm for 10 minutes, after which 100 μg of supernatant was loaded onto 50 μl streptavidin agarose resin (Myone C1 Dynabeads, Thermo Scientific) overnight at 4 °C. Resin-bound proteins were washed twice with RIPA lysis buffer, one with 1M KCl, once with 0.1 Na_2_CO_3_, once with 2M urea in 10 mM Tris-HCl, and three times with 50 mM ammonium bicarbonate, respectively. Proteins were reduced with 4 mM DTT for 25 minutes at 56 °C and alkylated with 8 mM iodoacetamide for 30 minutes at room temperature in the dark. Samples were digested with LysC (1:200 enzyme substrate ratio) and trypsin (1:100) overnight at 37 °C, after which the reaction was quenched with 2% FA. Peptides were desalted using Oasis HLB columns (Waters), dried *in vacuo* and stored at −80 °C until further analysis.

### LC-MS/MS

The enriched samples were analyzed with an UHPLC 1290 system (Agilent technologies) coupled to an Orbitrap Q Exactive HF X mass spectrometer (Thermo Scientific). Before separation peptides were first trapped (Dr Maisch Reprosil C18, 3 μm, 2 cm x 100 μm) and then separated on an analytical column (Agilent Poroshell EC-C18, 2.7 μm, 50 cm x 75 μm). Trapping was performed for 5 min in solvent A (0.1% FA) and eluted with following gradient: 0 - 13% solvent B (0.1% FA in ACN) in 10s, 13 - 44% in 95 min, 44 - 100% in 3 min, and finally 100 % for 1 min. Flow was passively split to 300 nl/min. The mass spectrometer was operated in data-dependent mode. At a resolution of 35.000 *m/z* at 400 *m/z*, MS full scan spectra were acquired from *m/z* 375–1600 after accumulation to a target value of 3e^6^ with a maximum injection time of 20 ms. Up to 15 most intense precursor ions were selected for HCD fragmentation at a normalised collision energy of 27%, after the accumulation to a target value of 1e^5^. MS/MS was acquired at a resolution of 30,000, with an exclusion duration 16s. Charge state screening was enabled, and precursors with an unknown charge state or a charge state of 1 were excluded.

### Data analysis

The raw data were analyzed using MaxQuant (version 1.6.3.4) for the identification and quantification of peptides and proteins ^19^. Data were searched against a database containing SwissProt Human proteome (downloaded 10/2018). Variable modifications were methionine oxidation, protein N-terminus acetylation and biotinylation by biotin-phenol on tyrosine (C_18_H_23_N_3_O_3_S). A fixed modification was cysteine carbamidomethylation. The first search was performed with a mass accuracy of ± 20 ppm and the main search was performed with a mass accuracy of ± 4.5 ppm. A maximum of 5 modifications and 2 missed cleavages were allowed per peptide. The maximum charge was set to 77+. For MS/MS matching, the mass tolerance was set to 0.5 Da and the top 8 peaks per 100 Da were analyzed. MS/MS matching was allowed for higher charge states, water and ammonia loss. The false discovery rate was set to 0.01. The minimum peptide length was 7 amino acids. Match between runs was performed with a time window of 0.7 minutes. Quantification was done label free with the MaxQuant algorithm with minimal ratio label count 2 and including unique and razor peptides. Further analyses were performed using Perseus version 1.6.2.2 ^20^, and Cytoscape ^21^ utilizing the GeneMANIA plugin ^22^.

### Immunoblotting

Cell lysates were incubated with Quickstain Cy5 protein-dye (GE Healthcare Bio-Sciences) for 30 minutes at room temperature, after which they were denatured and reduced in XT Sample buffer (Bio-rad) with 25 mM DTT at 95°C for 5 minutes. Proteins were separated on a 12% SDS-PAGE gel (Bio-rad) and electroblotted onto nitrocellulose membranes. Membranes were blocked in 5% non-fat milk in TBS with 0.1% Tween20 (TBS-T) and incubated with the following antibodies: α-pEGFR Y1068 (1:500, Abcam), Streptavidin conjugate Alexafluor-488 (1:10,000, Invitrogen), α-rabbit horseradish peroxidase conjugate (1:2000, Dako). When multiple antibodies of the same origin were used, membranes were stripped using Restore PLUS Western Blot stripping buffer (Pierce) to avoid cross-contamination of the secondary antibody. Detection was performed by enhanced chemiluminescent substrate (Pierce), or via fluorescence detection with the Amersham Imager 600 (GE Healthcare Bio-Sciences). Quantification was performed with Amersham imager analysis software (GE Healthcare Bio-sciences).

### Immunofluorescence and confocal microscopy

Cells were plated onto poly-L-lysine coated glass 18 mm coverslips and cultured as normal. They were fixed with 4% paraformaldehyde / 4% sucrose in phosphate-buffered saline (PBS) for 10 minutes at room temperature. Fixed cells were blocked with 10% normal goat serum in PBS for 30 minutes at 37 °C, and stained with following antibodies: α-EGFR (1:50, Cell Signaling), α-rabbit Alexa fluor 488 (1:1000, Invitrogen), and Streptavidin conjugate Alexafluor-594 (1:2000, Invitrogen). Confocal images were taken with a Zeiss LSM 710 with 63x 1.40 oil objective. Images consist of a z-stack of 7-9 planes at 0.39 μm interval, and maximum intensity projections were generated for analysis and display.

## Results & Discussion

### EGFR-APEX2 is functional, specific, and localizes correctly

We first characterized the functionality of the newly fused EGFR-APEX2 fusion protein, where APEX2 was incorporated at the C-terminus of the EGFR. As a control, the well-characterized EGFR-eGFP control was used. No difference in expression levels between the two constructs could be observed (Figure S1A). First, we sought to assess the functionality of the APEX2 enzyme in the newly formed fusion protein. To address this, EGFR-APEX2 or EGFR-eGFP was introduced into HEK293T cells, and the activity of the enzyme was tested in presence of biotin phenol and H_2_O_2_. Western blot analysis of biotinylated proteins shows that EGFR-APEX2, and not the negative control constructs, causes biotinylation on proteins in a wide range of molecular weights. Moreover, biotinylation was exclusively observed in conditions where cells were pre-incubated with substrate (biotin-phenol) and activated with H_2_O_2_ (Figure 1A). Second, we assessed whether replacement of eGFP to APEX2 did not alter the functionality of the EGFR receptor. To this end, we placed the cells in starvation media without fetal bovine serum for 30 to 90 minutes and stimulated EGFR with EGF for 30 minutes. We then analyzed autophosphorylation of the receptor on tyrosine 1068 using western blot analysis and saw a clear increase in autophosphorylation after stimulation (Figure S1B), indicating that EGFR can still be activated in the presence of APEX2 at its cytoplasmic tail. We then evaluated EGFR activation following EGF and TGF-α stimulation over time. We observed that while TGF-α-induced EGFR activation causes a sustained increase in EGFR phosphorylation, EGF-induced EGFR phosphorylation decreases over time (Figure S1C). Finally, we performed immunofluorescence staining experiments to visualize the localization of our fusion protein in HEK293T cells. We found that EGFR-eGFP localizes predominantly to the plasma membrane and that a smaller pool of receptors can be distinguished in the endoplasmic reticulum (Figure 1B). We observed very similar localization patterns for our EGFR-APEX2 fusion protein via visualization of biotinylated proteins, selectively in conditions where we performed the APEX2-driven biotinylation reaction (Figure 1C). Taken together, we concluded that EGFR-APEX2 was functional and suitable for further biological experiments.

**Figure 1.**
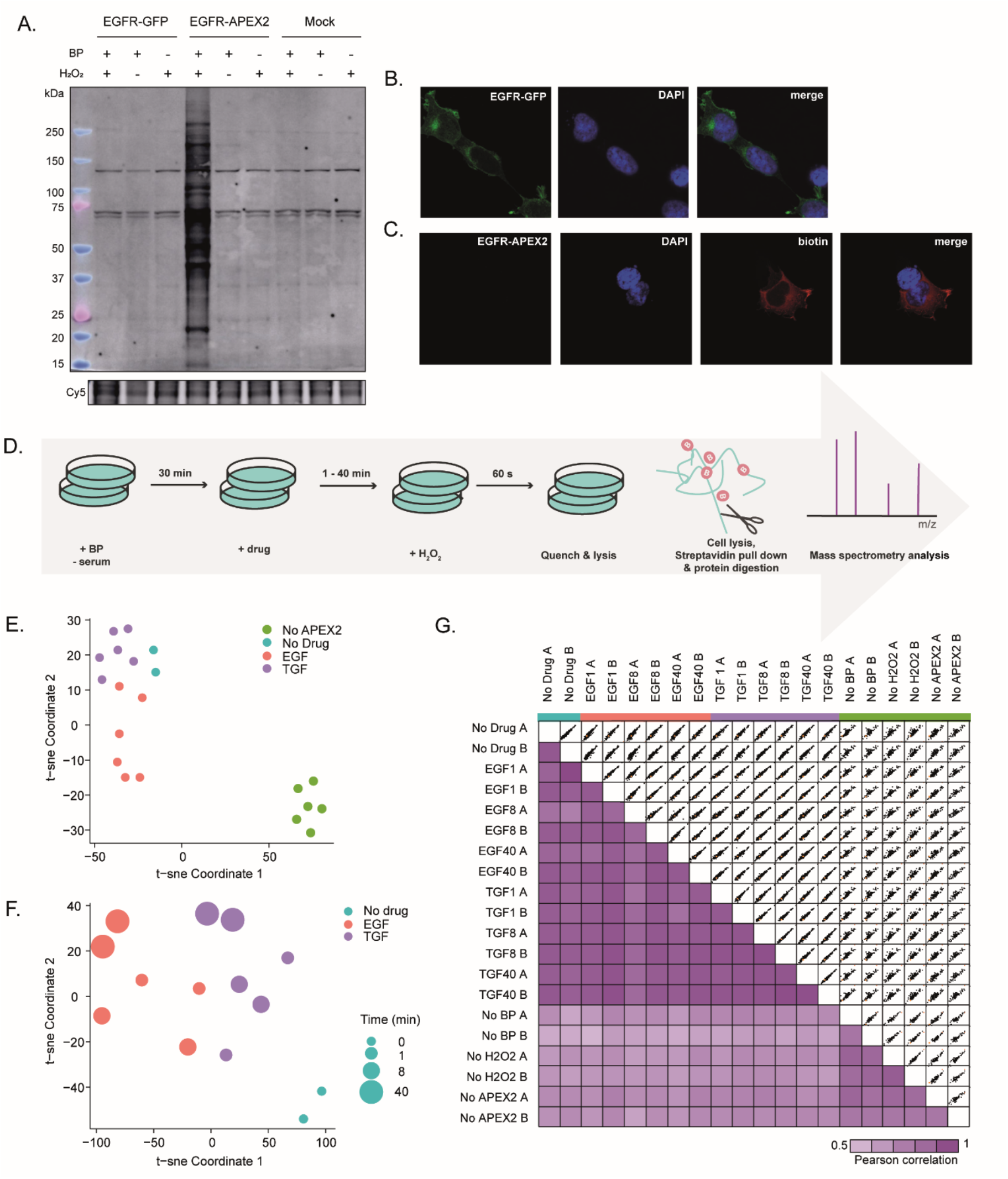
EGFR-APEX2 activation, localization, and data quality. (A) Western blot of biotinylated proteins in different ‘APEX2’ and ‘No APEX2’ conditions. Biotinylation only occurred with the APEX2 fusion construct present, after incubation with BP and subsequent stimulation with H_2_O_2_. Cy5 dye indicates protein loading. (B) Localization of the EGFR-GFP control construct. Expression is predominantly localized to the plasma membrane and ER. Cy5 dye indicates protein loading. (C) EGFR-APEX2 is localized similarly to the EGFR-GFP control construct, as the biotinylation pattern matches EGFR localization. (D) Experimental workflow and data quality. HEK293T cells were transfected with EGFR-APEX2 construct and incubated for 48 hours before the start of the experiment. At the start of the experiment, cell culture media was depleted from serum and supplemented with biotin phenol (BP). After 30 minutes, EGF or TGF-α was added. The APEX2 reaction was initiated by addition of H_2_O_2_, and quenched after 60 seconds, after which the cells were lysed. Biotinylated proteins were extracted from the cell lysate using streptavidin-coated beads, digested, and measured by mass spectrometry. (E,F) T-sne plot of all experimental conditions reveals a clear separation between ‘APEX2’ and ‘No APEX2’ conditions, as well as a defined separation of EGF and TGF-α treated samples over time, respectively. Pearson correlation plot displaying a high correlation between all ‘APEX2’ conditions, while ‘no APEX2’ and ‘APEX2’ conditions correlate significantly less. N=2 per condition.

### Enrichment of biotinylated proteins leads to enrichment of proteins related to EGFR signaling

Next, we sought to follow EGFR interactome dynamics upon stimulation with EGF and TGF-α. We therefore incubated HEK293T cells expressing EGFR-APEX2 with biotin-phenol (BP) in the absence of serum for 30 minutes, after which we stimulated with either EGF or TGF-α (100 ng/ml) for 1, 8, or 40 minutes. Then, we made snapshots of the EGFR interactome by addition of H_2_O_2_ for 60 seconds, after which the reaction was quenched and cell pellets were collected. An equal amount of cell lysate was loaded onto streptavidin-coated beads and incubated overnight to capture biotinylated proteins (Figure S2A). After stringent washing steps, biotinylated proteins were digested off-bead and analyzed using LC-MS/MS (Figure 1D). Next to the two drug treatments, we included several ‘no APEX2’ negative control conditions, where we systematically excluded BP incubation, H_2_O_2_ stimulation, or both (Figure S2B).

Visualization of all identified and quantified proteins (2,517) in a tSNE plot showed a clear separation between ‘APEX2’ and ‘no APEX2’ experimental conditions, as well as a separation between EGF, TGF-α, and unstimulated samples (Figure 2E,F). Additionally, we observed a high Pearson correlation between all ‘APEX2’ conditions, while ‘no APEX2’ and ‘APEX2’ conditions correlated significantly less (Figure 1G), confirming the selectivity and reproducibility of the biotin-based enrichment process.

**Figure 2.**
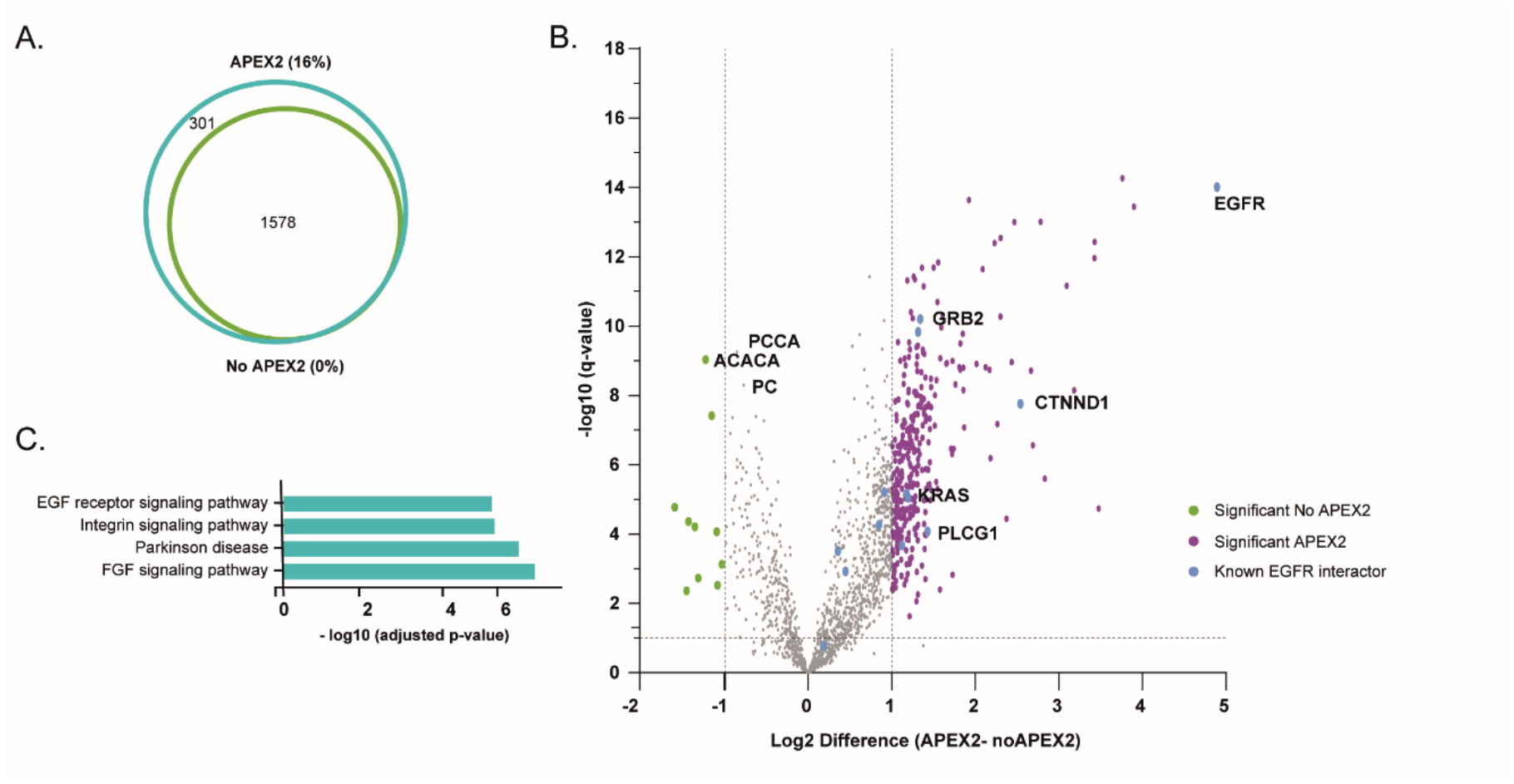
(A) Venn diagram illustrating the overlap in identified proteins in the ‘APEX2’ versus ‘No APEX2’ conditions. (B) Volcano plot of all ‘APEX2’ proteins versus ‘No APEX2’ proteins, where a clear skewed distribution of protein intensities and significance towards the ‘APEX2’ side can be observed, indicating the success of the enrichment strategy. (C) GO overrepresentation analysis showed a clear enrichment of EGFR-related biological processes amongst ‘APEX2’ proteins.

Next, we filtered based on proteins that were identified in less than three conditions in either ‘APEX2’ or ‘no APEX2’ groups, resulting in more than 300 proteins uniquely identified in the ‘APEX2’ conditions (Figure 2A). Then, we tested which proteins had a significantly higher intensity profile in ‘APEX2’ versus the ‘no APEX2’ conditions (Figure 2B) and selected these proteins (fold change >2 and q-value <0.05) for further analysis, together with the ‘APEX2’ unique proteins, resulting in a total of 442 proteins (Table S1). Among these proteins were many known EGFR interactors with significantly higher intensity in the ‘APEX2’ conditions, including EGFR itself, and the well-known EGFR signaling cascade proteins GRB2, and PLCG1, that are adaptor and second messenger proteins in the activation of the ras, and PKC signaling pathways, respectively. Moreover, three endogenously biotinylated proteins (PCCA, ACACA, and PC) were much more intense in the ‘no APEX2’ conditions and therefore filtered out. Statistical overrepresentation analysis of Reactome pathways of the total pool of identified EGFR interactors confirmed a clear enrichment of EGF receptor and related signaling cascades (Figure 2C).

### Phosphorylation of EGFR interactors

Since it is well known that EGFR signaling is highly dependent on tyrosine kinase activity, we examined to which extent we could see this reflected in our EGFR interactome datasets. To understand the extent of phosphorylation dynamics, we searched the RAW files for STY phosphorylated peptides and normalized the intensities of identified phosphosites to the relative protein ratio at each time point. Strikingly, we found that in our entire proteome dataset we identified 243 class I phosphosites (e.g. with a localization probability score higher than 0.75 and identified in at least two APEX2 replicates), corresponding to 132 unique proteins (Table S1). If we consider the whole quantified proteome dataset (1,879 proteins), this results in 12.9% of our proteins being phosphorylated, all without specific phosphopeptide enrichment.

#### EGFR phosphorylation sites

Of our identified phosphosites, 41 were found to be significantly more abundant in the ‘APEX2’ conditions vs ‘no APEX2’ conditions, among which we identify three EGFR phosphorylation sites; S1166, Y1197, and T693. T693 (also often referred to as T669) is the major EGFR activation site after EGF stimulation, although it has also been shown to be constitutively active ^23^. Phosphorylation is performed by p38 MAP kinase ^24^. Y1197 is also known to be phosphorylated by MAP kinases upon EGF stimulation. S1166, as well as other serine and threonine phosphorylation sites located in the cytoplasmic tail of EGFR between amino acids 1047-1072, was shown to be phosphorylated by PKA, thereby positively regulation tyrosine kinase activity ^25^. Altogether these results show that our EGFR-APEX2 approach is very efficient in monitoring EGFR activation, as all of these indicative phosphorylation sites have an over five times higher intensity in the ‘APEX2’ versus ‘non-APEX2’ conditions. The lack of identification of another major EGFR activation site, Y1068, which is routinely used to monitor EGFR activation in low throughput studies such as western blot analysis, can be explained by the utilized proteomics workflow. During sample preparation proteins were digested with Trypsin, with a high cleavage specificity for Lysine (K) and Arginine (R) on the N-terminal side, as indicated with the purple horizontal lines in Figure S3A. Since, in the following database search, typically only peptides with a length of maximum 25 amino acids are allowed, the peptide which contains the activation site (previously referred to Y1068 but actually located at Y1092 in the Uniprot EGFR sequence, indicated in the orange square) was not identified as it consists of 31 amino acids. However, searching with an extended amino acid range did indeed result in the confident identification and localization of the Y1092 activation site (Figure S3B).

#### Phosphorylation dynamics upon EGFR stimulation

To identify which of these APEX2-enriched phosphorylation sites are biologically relevant in EGFR signaling dynamics, we further selected only those phosphorylation sites that showed a clear difference (at least two-fold) between EGF and TGF-α in at least one time point, or that showed a clear increase or decrease (at least two-fold) between either EGF and/or TGF-α compared to the ‘No Drug’ control situation. The resulting 26 phosphorylation sites are displayed in Table 1.

**Table 1.** Significantly regulated phosphorylation sites in ‘APEX2’ upon EGFR stimulation. Phosphorylation sites with a log2 fold change >1 between EGF and TGF-α in at least one time point, or between EGF and/or TGF-α compared to control are included. Values represent log2 intensity differences between conditions.

| Phosphosite | EGF 1min/ND | TGF-α 1min/ND | EGF/TGF-α 1 min | EGF/TGF-α 8 min | EGF/TGF-α 40 min |
| --- | --- | --- | --- | --- | --- |
| EIF5B_S66 | 1.97 | 0.99 | 0.98 | 0.07 | 0.36 |
| BCLAF1_S512 | -1.62 | -0.73 | -0.89 | 0.77 | 0.85 |
| CHMP7_T391 | 0.74 | 0.49 | 0.24 | -0.68 | -1.69 |
| CCDC38_S127 | 1.05 | 0.76 | 0.29 | -0.56 | 0.06 |
| NOP2_S732 | 1.12 | 0.12 | 1.00 | 0.52 | 1.10 |
| TCOF1_S1228 | 0.38 | -0.16 | 0.54 | 0.12 | 0.48 |
| DDX21_S121 | 0.45 | 0.02 | 0.43 | 3.03 | 1.15 |
| RRP1B_S245 | 1.19 | 0.96 | 0.23 | 0.47 | 0.72 |
| NAV2_S503 | 2.16 | -0.55 | 2.71 | -1.58 | 1.78 |
| TCOF1_S171 | -0.34 | 0.35 | -0.69 | 1.05 | 3.08 |
| PER1_T164 | -0.71 | -1.70 | 0.99 | -0.52 | 0.48 |
| TOP2A_S1393 | 2.09 | 0.70 | 1.39 | -0.69 | 0.23 |
| SMARCA5_S66 | 1.59 | 1.17 | 0.42 | 0.94 | -0.42 |
| CHD4_S515 | 0.69 | 0.55 | 0.14 | -0.54 | 1.41 |
| PER1_Y167 | -0.71 | -1.73 | 1.02 | -0.52 | 0.14 |
| RRP1B_S732 | -0.46 | -1.01 | 0.55 | -2.42 | 1.42 |
| NOP56_S520 | 0.05 | -0.16 | 0.21 | 1.78 | 1.53 |
| ZMYM4_S1284 | 1.83 | 0.41 | 1.41 | 1.71 | 0.41 |
| ZBTB40_S223 | 1.47 | 0.51 | 0.97 | 0.31 | 1.34 |
| ECT2_T359 | 1.83 | 1.38 | 0.44 | 0.23 | 1.84 |
| EIF4B_S459 | -1.18 | -1.34 | 0.16 | -2.61 | -1.63 |
| MLLT4_S1275 | 0.31 | -0.39 | 0.69 | 1.62 | 2.13 |
| ABI1_S225 | 0.89 | -0.99 | 1.89 | 1.54 | 2.49 |
| EGFR_T693 | 1.82 | 1.71 | 0.11 | 0.30 | 0.55 |
| DNAH7_S1252 | 2.49 | 0.22 | 2.27 | 1.73 | 1.91 |
| EGFR_Y1197 | 0.67 | 0.19 | 0.48 | -0.15 | 1.54 |

Higher in TGF $\alpha$ Higher in EGF
log<sub>2</sub> (EGF/TGF $\alpha$ )

We found two translation initiation factors with phosphorylation sites showing striking dynamical behavior over the time course and between the two treatment conditions. The S66 phosphorylation site of Eif5b is a known indicator of translation initiation, which, in our data, displayed very high initial activation in both EGF and TGF-α conditions, although with a much higher response in EGF-stimulated cells. This initial activation was followed by rapid de-phosphorylation at 8 minutes, which in turn was followed by re-phosphorylation at the latest time point, in both EGF and TGF-α treated cells. EIF4B displayed de-phosphorylation of S459, suggesting that de-phosphorylation of this site plays a role in the initiation of translation. Interestingly, EIF4B remained dephosphorylated in EGF stimulated conditions over time, but conversely became phosphorylated in TGF-α conditions at later time points. Bclaf1 is a transcriptional repressor of survival genes. We observe rapid dephosphorylation in both EGF and TGF-α conditions, which hints towards activation of survival genes. Interestingly, rephosphorylation at later time points is more pronounced in TGF-α than EGF conditions.

### Temporal behavior of hallmark proteins

We next sought to identify differences in EGFR interactome dynamics between the two different stimulus conditions. We therefore investigated the behavior of three hallmark vesicle proteins over time. Rab5 is a marker of early endosomes. As internalization and early endosome encapsulation of EGFR is a common trafficking route after both EGF and TGF-α-induced EGFR activation, no bias in interaction levels is expected between the two different treatments. Indeed, the relative amount of RAB5 in close proximity to EGFR increases after both stimuli, and stabilizes after 8 minutes (Figure 3A, upper panel). As a marker of late endosomes, RAB7A is a validated indicator of protein degradation. Characteristically, the switch from RAB5 to RAB7A defines the evolution of early to late endosomes. Interestingly, we observe an initial decrease in RAB7A in the EGF treated samples compared to the TGF-α samples after 8 minutes of stimulation, which normalizes at the 40-minute time point (Figure 3A, middle panel). This unexpected difference in RAB7A localization pattern could not be explained by potential modification of the RAB7A peptides used for quantification; previous literature has shown that Rab7a undergoes multiple modifications (ubiquitination and phosphorylation) specifically in the early time points ^3^. However, quantification of RAB7A in the current dataset was found not to be based on the peptides carrying earlier reported modifications. Finally, RAB11 is a marker of recycling endosomes, and can therefore be used as an example for TGF-α -mediated EGFR recycling, as the majority of EGFR recycles back to the plasma membrane upon stimulation (91% for TGF-α versus 22% in EGF-stimulated samples, respectively ^3^). In our data, we can indeed see a clear increase in proximity upon stimulation of the EGFR in the initial phase, which continues to increase over time for the TGF-α treated samples. In the EGF conditions however, we lose detection of RAB11 after 8 minutes, indicating that the amount of EGFR that is enclosed in RAB11 coated vesicles is below detection level (Figure 3A, lower panel). Taken together, these data clearly highlight the efficacy of the APEX2 proximity labeling protocol in deducing protein cellular location with the help of so called ‘bystander’ proteins.

**Figure 3.**
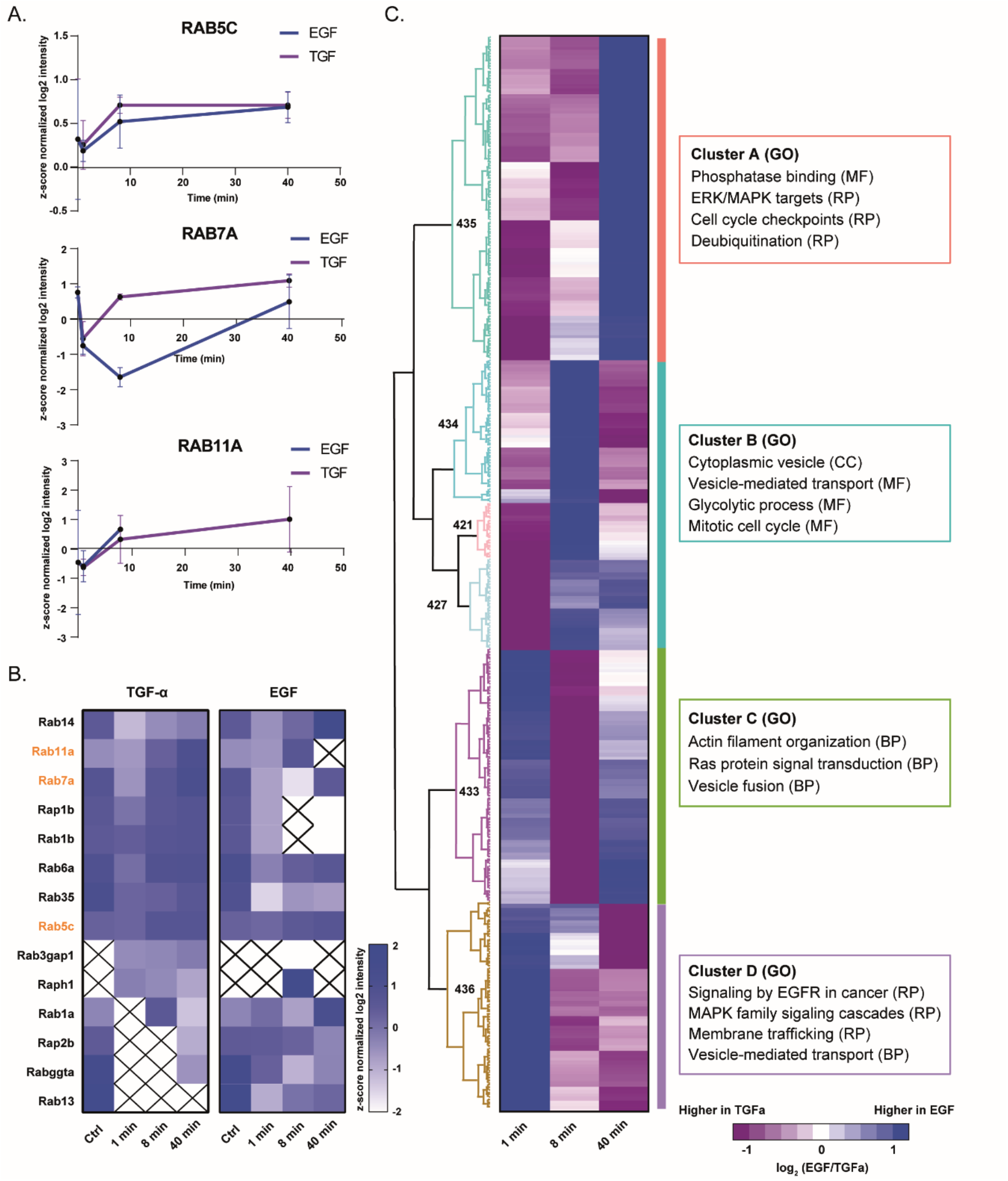
Stimulus-induced EGFR interactome perturbation. (A) Z-score normalized intensity plots of hallmark endosomal proteins following EGF and TGF-α stimulation, respectively, indicating the relative subcellular location of EGFR over different time points. (B) Temporal behavior of Rab family proteins yield information on vesicle-specificity and function. X – protein not identified at the specified time point. (C) Heatmap illustrating the differences in z-score normalized protein intensity between EGF and TGF-α stimulated samples for each time point. Hierarchical clustering based on Euclidean distance of all ‘APEX2’ proteins revealed four main clusters of proteins.

#### Temporal behavior of other Rab and rap family proteins

Next to these well-established and characteristic Rab proteins, we identify several other Rab proteins. The Rab family of small GTPases are key players of intracellular membrane trafficking, regulating the formation of transport vesicles to their fusion with membranes ^26^. Comparing the relative intensity of several, lesser studied members of the Rab family of proteins, as well as Ras-related GTP-binding proteins (Rap), between differential EGFR trafficking conditions, could yield relevant information on the function of these proteins. We therefore plotted the temporal behavior of all Rab and Rap proteins following TGF-α, and EGF stimulation, respectively (Figure 3B).

RAB1B and RAP1B intensities increase over time for TGF-α samples, while they are not detected, or very low abundant in the later time points for EGF-stimulated samples. As RAB1B recruitment to the Golgi is known to enhance vesicle secretion ^27^, it seems plausible that it facilitates receptor recycling towards the plasma membrane. RAP1B receptor binding promotes GABA receptor surface expression by facilitating receptor recycling ^28^, a pattern that fits TGF-α-induced EGFR trafficking. Other interesting observations include RAB3GAP1 and RAPH1, which are detected only after EGFR ligand activation. More specifically, both are absent in the control condition and readily detected after TGF-α addition followed by a steady increase over time. Conversely, both RAB3GAP1 and RAPH1 are detected after EGF stimulation only at the 8-minute time point, displaying a large difference in intensity. RAB3GAP1 is the catalytic subunit of a GTPase activating protein with a high specificity for the Rab3 subfamily and converts active RAB3-GTP into its inactive form, RAB3-GDP. In synaptic vesicles, this inactivation of RAB3 results in clathrin-mediated endocytosis and recycling of these vesicles ^29^. RAPH1 is a known mediator of localized membrane signals, and a known interactor of SRC and ABL1, and might therefore be implicated in early receptor endocytosis and cytoskeleton reorganization. Hence, both of these proteins have a likely role in the early stages of receptor endocytosis, a process that is more dominant and stable after TGF-α stimulation.

On the contrary, RAB13 seems to be more EGF-specific, as it is not detected upon TGF-α stimulation, while it increases steadily over time in the EGF conditions. Furthermore, RAP2B and RABGGTA are highly abundant quickly after EGF receptor activation, and slowly decrease in intensity over time. After TGF-α stimulation, however, RAP2B is not detected in the earliest two time points. This is unexpected, as several reports describe Rap2b to be recruited by activated EGFR, thereby activating PLC-ε signaling and further downstream processes, although these data seem to be based on EGF-stimulated EGFR only ^30,31^. RABGGTA is a geranylgeranyl transferase and catalyzes the transfer of a geranylgeranyl lipid moiety to the c-terminus of certain Rab proteins, thereby anchoring them to their target membrane ^32,33^. Which membrane, however, remains unknown, and our data indicates that it might be preferentially involved in degradation-related vesicle-membranes. Overall, these data provide a good indication of the role of several lesser-studied vesicle-related proteins and give a first glance at their specificity towards different vesicle-types. Further validation studies are required for detailed characterization.

### EGFR interactome significantly changes upon stimulation

To investigate the EGFR dynamic interactome over time between the different stimulus conditions, we first normalized the protein intensity values to their non-stimulated control intensities. Next, we calculated the differences in protein intensity between EGF and TGF-α stimulated samples for each time point and performed Euclidean distance hierarchical clustering of all ‘APEX2’ proteins. In the resulting heatmap (Figure 3C) we can distinguish the relative abundance of EGFR interacting proteins in both the EGF and TGF-α conditions at the different time points, as the color scheme indicates the difference in intensity between both treatments. We can distinguish four main protein clusters, each with a distinct bias towards one of the ligands at one or more of the investigated time points. To get an indication of which proteins were noticeably biased towards one of the two signaling routes (i.e. EGF and TGF-α), we performed GO overrepresentation analyses on cellular component (CC), biological function (BF), molecular function (MF), and Reactome Pathways (RP).

Cluster A contains proteins that are predominantly proximal to EGFR upon TGF-α stimulation at the early time points but shift towards EGF at the later time point. This was confirmed by GO enrichment analyses, where clear EGF-related terms are enriched, such as the dominant presence of ERK/MAPK targets. Moreover, EGF-related EGFR signaling termination was observed, as indicated by an enrichment in protein phosphatase 2 (PP2A) subunits. Activation of PP2A results in downregulation of both PI3K and MAPK pathways ^34^, both of which are classical EGF-related signaling pathways. Conversely, cluster D switches from EGF to TGF-α from 1 minute to 40 minutes and is therefore enriched in proteins from EGF-directed signaling pathways, including MAPK family signaling cascades. Also, proteins involved in vesicle-mediated transport and membrane trafficking are enriched, indicating TGF-α-characteristic receptor recycling and prolonged activation. Clusters B and C show a more dynamic behavior, where cluster B shifts from a bias in TGF-α at 1 minute to EGF after 8 minutes followed by a almost equal representation after 40 minutes, indicating that these proteins are active after EGFR stimulation with both TGF-α and EGF. Not surprisingly, overrepresented GO terms include general cellular processes such as cell cycle and glycolytic processes, and general vesicle-mediated transport. Cluster C displays the opposite behavior at 1 and 8 minutes, and contains proteins involved in RAS protein signal transduction, as well as the more general terms actin assembly and reorganization, and vesicle fusion.

### Deducing cellular localization and biological processes from bystander proteins

Based on previous research, we expect the biggest alterations in EGFR interactome at the 40-minute time point, where we can start observing differences in receptor trafficking (*e.g.* degradation versus recycling) ^3^. These differences in receptor distribution are expected to result in vastly different proteins in the proximity of EGFR, while more subtle differences in receptor stimulation and activation are expected to be less obvious in the study of the interactome. Hence, we further examined the proteins of clusters A and D and performed a functional network analysis on both clusters, as visualized in Figures 4A and 4B, using the STRING database in combination with GO enrichment analyses. GO terms with FDR-corrected p-values lower than 0.01 were considered for network mapping.

**Figure 4.**
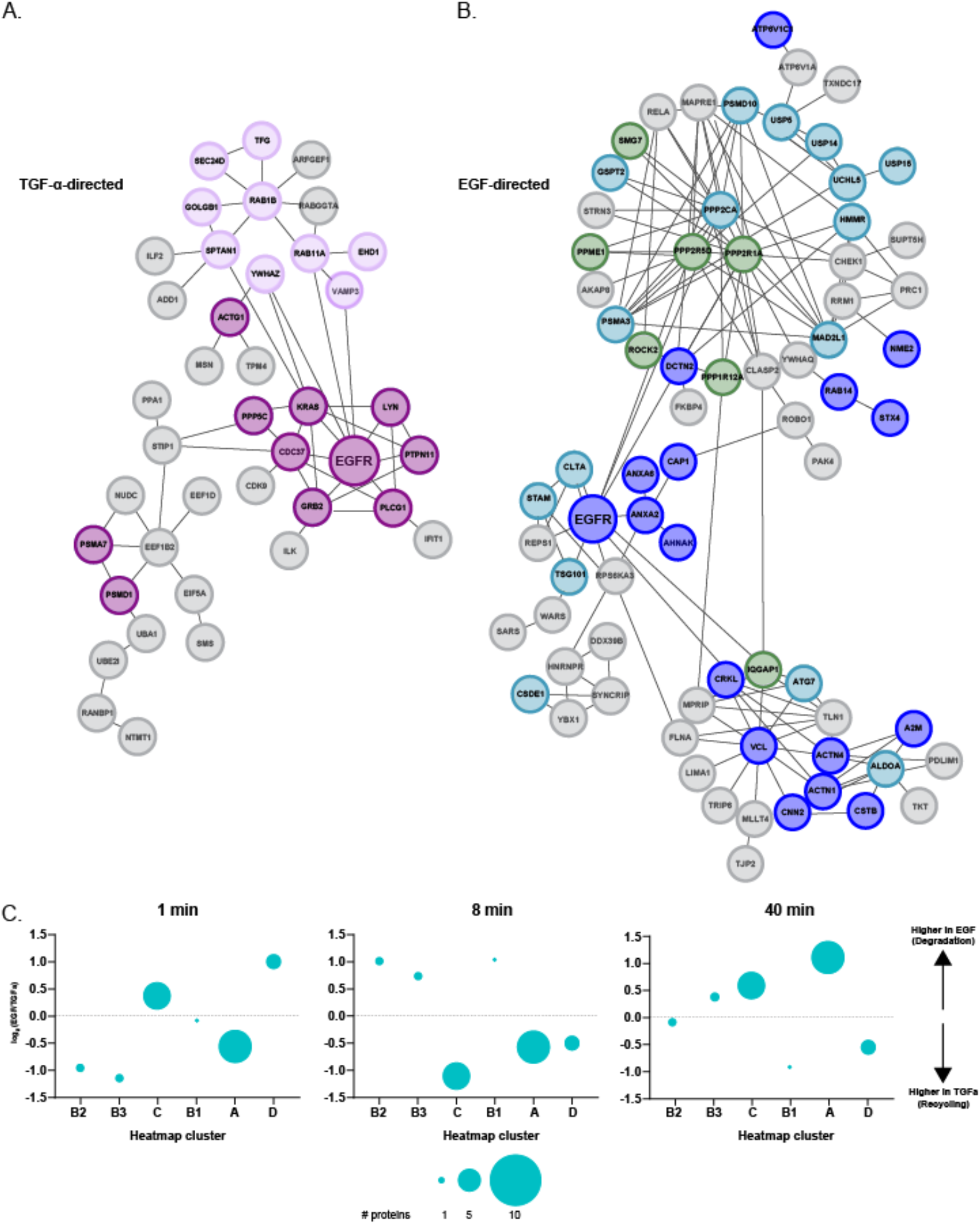
Protein interaction networks from heatmap clusters that are (A) TGF-α-directed, and (B) EGF-directed at the 40-minute time point. Colors indicate membership to sub clusters, as defined by enrichment in GO terms. lilac – recycling endosomes, purple – membrane interactors, dark blue – vesicle transport, light blue – protein degradation, green – phosphatases and other receptor-deactivating proteins. (C) Differential clustering of vesicle proteins reveals differences in timing between EGF and TGF-α signaling duration. While early signaling (1 minute, left panel) shows an equal distribution of vesicle proteins between EGF and TGF-α, differences are observed in the later time points. At the 8-minute time point, vesicle proteins are more dominantly present in the TGF-α stimulated samples, indicating that at that time point, EGFR is present in recycling vesicles (middle panel). At the late time point (40 minutes) however, we observe a shift towards EGF, and therefore towards degradation-related vesicles (right panel).

Figure 4A displays the interaction network of cluster D, biased towards TGF-α at the 40 min time point. We included 85 nodes, with a total of 68 edges, using a confidence cut off score of 0.6. Of these, 43 nodes were excluded for further network analysis because they were not connected to the major network. GO enrichment analyses revealed two main functional sub clusters, highlighting the presence of EGFR near recycling endosomes (green) and membrane interactors (blue), respectively. This indicates that the receptor is recycled and active, as we here identify a number of tyrosine kinases and phosphatases that are known to signal downstream of EGFR, such as tyrosine-protein kinase Lyn (LYN) tyrosine-protein phosphatase non-receptor type 11 (PTPN11). Membrane interaction is further supported by the presence of the well-known EGFR membrane interactor GRB2, and Moesin (MSN), where the latter functions as an anchor between microtubules and the plasma membrane ^35^. Moreover, the presence of cyclin-dependent kinase CDK9, and the Hsp90 co-chaperone CDC37, a protein that can stabilize protein kinases via HSP90 interactions, further strengthen the enrichment of membrane interacting proteins.

Next, we investigated some smaller protein networks that were not directly connected to the major EGFR network, however, we believe are associated to EGFR function. First, a small complex of RAP2B and SERPINB6. RAP2B, as mentioned before, is recruited by activated EGFR and was marked as a membrane interactor. According to Reactome data, SERPIN6 is also a membrane interacting protein, and was shown to be downregulated in EGFR tyrosine kinase domain mutants ^36^, indicating that it is involved in early EGFR activation and endocytosis. A second network consists of NME1, POLA2, TXNRD1, and GLRX3. Nme1, also known as NM23, facilitates clathrin-dependent internalization of EGFR, since knockdown of NM23 reduced EGFR endocytosis ^37,38^. TXNRD1 is a key antioxidant enzyme, and inhibition of TXNRD1 has been shown to sensitize EGFR-related carcinomas to treatment ^39,40^. Stabilization of GLRX3, another antioxidant enzyme, increases EGFR expression in nasopharyngeal carcinoma cell lines ^41^. Taken together, our results show that several other TGF-α-directed EGFR proximity proteins can in fact be added to the functional EGFR repertoire.

To plot the EGF-biased interaction network derived from the proteins in cluster A, we supplemented the group of proteins with EGFR (as it is located in cluster D), resulting in the network depicted in Figure 4B. The computed protein network consisted of a total of 134 nodes, with 155 confident edges. Again, only nodes with confident edges between major nodes of the main interaction network were kept for further analysis, so that 64 nodes were discarded. Here, GO analyses indicated clear enrichment of vesicle transport (red), protein degradation (orange) and phosphatases and other proteins that deactivate receptor signaling (purple). While vesicle transport is a quite general GO term, closer inspection of the proteins that were included in the enrichment of this term hint towards a more specified type of vesicle transport. Both V-type proton ATPase subunit C 1 (ATP6V1C1) and its catalytic subunit A (ATP6V1A) indicate EGFRs proximity to low pH vesicles, such as lysosomes.

Other proteins from cluster A that are not connected to the major network but are likely still relevant for EGF-related EGFR interactions, are, among others, OTUD7B and USP24, both of which are deubiquitinating enzymes. EGFR is ubiquitinated when activated, which eventually leads to degradation of the receptor. Deubiquitinating enzymes however, can prolong receptor activation by removing ubiquitin chains ^42^. OTUD7B, also known as Cezanne-1, has been shown to enhance receptor signaling by stabilizing EGF-activated EGFR ^43^. USP24 is downregulated in EGFR adenocarcinomas, as well as in EGF-treated primary lung cells. In the same cells, knockdown of USP24 increases cell numbers and cell viability ^44^. Taken together, these proteins are interesting EGFR interactors that can play a role in downstream EGFR processes that involve cell proliferation and viability.

#### Differential clustering of vesicle proteins reveals information on timing and signaling duration

As multiple of our employed enrichment analyses clearly revealed vesicle-related proteins, we hypothesized that the localization of the different vesicle proteins at the different time points could contain information on the timing and duration of EGFR signaling. We therefore investigated the overlap of the proteins from the heatmap with the Exocarta Exosome database and found that 30 out of 441 proteins (6.8%) in the heatmap are in the Exocarta top 100 extracellular vesicle proteins. It should be noted, however, that these proteins are not exclusively found in extracellular vesicles but are considered to be identified more often in exosomes than in the general proteome. Of these, 22 are clearly EGF biased at the 40-minute time point as they can be found in cluster A.

We know that EGFR proximity proteins can reveal details on the location of the receptor via the so-called ‘bystander’ proteins. As activated EGFR spends the majority of its lifetime in intracellular vesicles, we hypothesized that we could use the data from all vesicle related Exocarta top 100 proteins to yield information on the timing and localization of the receptor between the different ligands. To this end, we plotted the average EGF/TGF-α intensity ratio from all vesicle proteins in each heatmap cluster (Figure 3C) at each time point after stimulation, depicted in Figure 4C. The data in the plots clearly illustrates differential timing between the two ligands. At the earliest time point, there is no difference in the average intensity between the vesicle proteins, illustrated by the equal spread of vesicle proteins across the intensity axis. A clear shift can be observed in subsequent time points, where the majority of vesicle proteins are TGF-α focused at 8 minutes, while at the ‘late’, 40-minute, time point there is a clear shift for these vesicle-related proteins towards EGF, corresponding to the degradation-related proteins in heatmap cluster A. In summary, these data indicate that upon stimulation of EGFR by EGF and TGF-α, receptor recycling is initiated early in the signaling process, whereas degradation is a slower process. This evidence is strengthened further by the behavior of Rab and Rap proteins from Figure 3B, where receptor endocytosis and recycling proteins were earlier and more dominantly expressed near EGFR after TGF-α stimulation, compared to EGF stimulation.

## Conclusion

In conclusion, we showed that with a label free APEX2 live cell proximity method we were able to map EGFR subcellular location at different time points after stimulation. Utilizing the fast and concise biotinylation of proximity proteins by APEX2, we were able to detect slight differences in early signaling kinetics between TGF-α and EGF, and identified ligand-specific vesicle-related proteins, thereby increasing our knowledge on RTK signaling and differential trafficking.

## Acknowledgements

This research was part of the Netherlands X-omics Initiative and partially funded by NWO, project 184.034.019 and the Horizon 2020 program INFRAIA project Epic-XS (Project 823839). We would like to thank Bohui Li for help with the generation of the t-sne plots.

## Data availability

The mass spectrometry proteomics data have been deposited to the ProteomeXchange Consortium via the PRIDE partner repository (http://www.ebi.ac.uk/pride/archive/) with the data set identifier PXD024136.

## Supplemental figures

**Figure S1.**
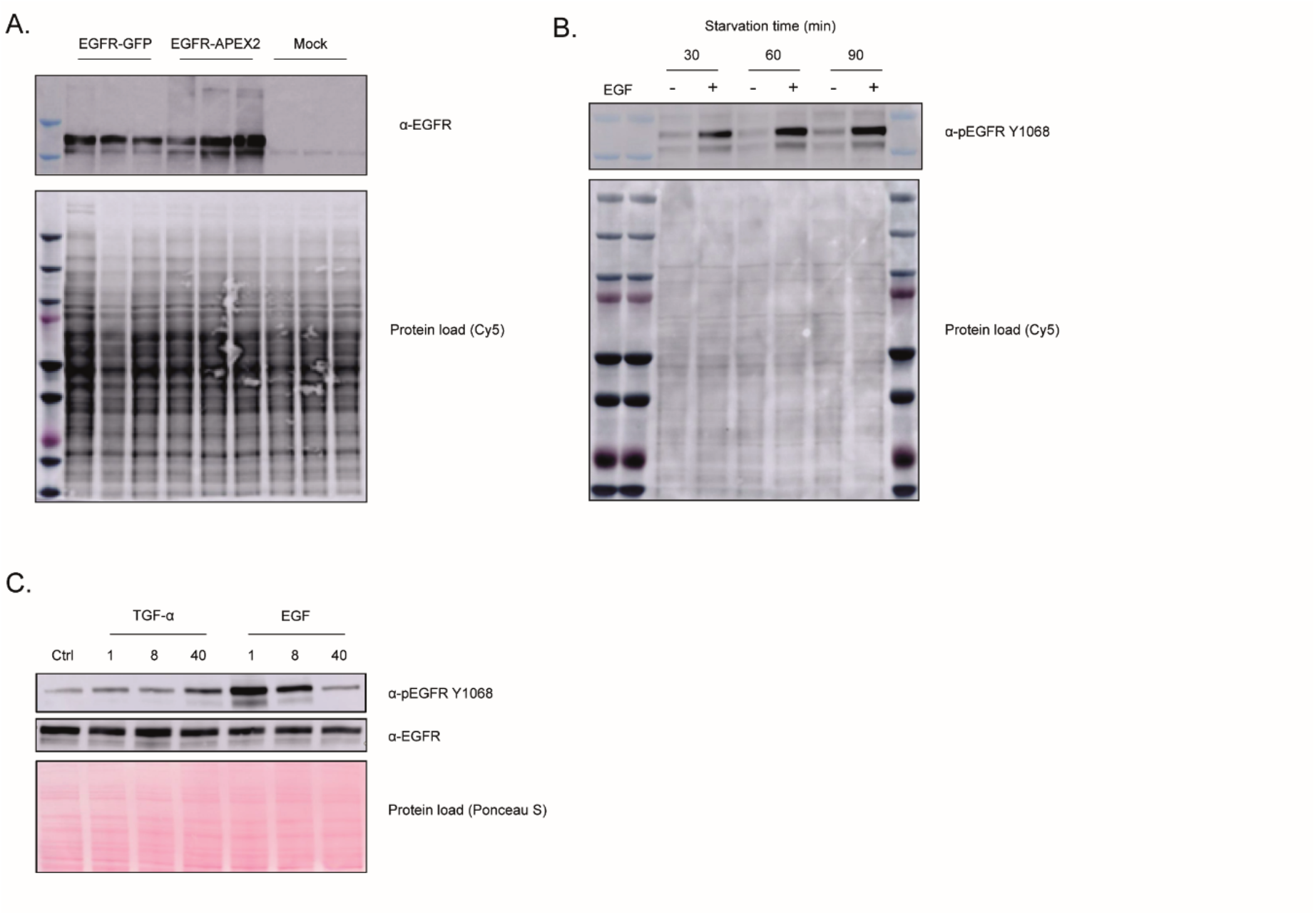
Validation of EGFR-APEX2 expression and activation in HEK293T cells. (A) Western blot evaluation of EGFR expression for EGFR-GFP and EGFR-APEX2 constructs. (B) Western blot of EGFR Y1068 phosphorylation upon EGF stimulation, following different periods of serum starvation. (C) Evaluation of EGFR activation following EGF and TGF-α stimulation over time. While TGF-α-induced EGFR activation causes a sustained increase in EGFR phosphorylation, EGF-induced EGFR phosphorylation decreases over time.

**Figure S2.**
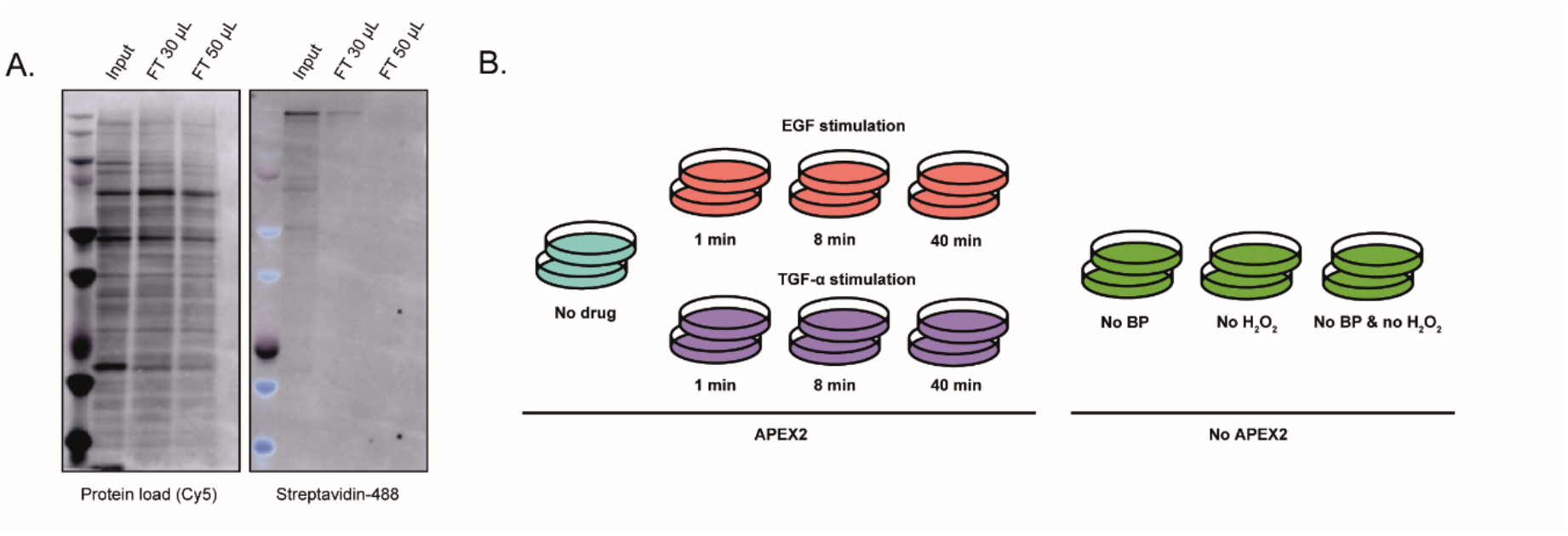
Biotin enrichment efficacy and workflow of the APEX2 proximity labeling experiment. (A) Western blot analysis of enrichment efficacy using different amount of streptavidin-coated beads. (B) Experimental conditions utilized in this study. Duplicate biological replicates were prepared for each experimental condition. Samples belonging to the ‘APEX2’ conditions were both incubated with BP and stimulated using H2O2, as described in the experimental workflow, and stimulated with either EGF or TGF-α for 1-40 minutes, respectively. The ‘No APEX2’ condition comprised of samples that were deprived from one or more of the APEX2 reagents, being BP, H2O2, or both, respectively.

**Figure S3.**
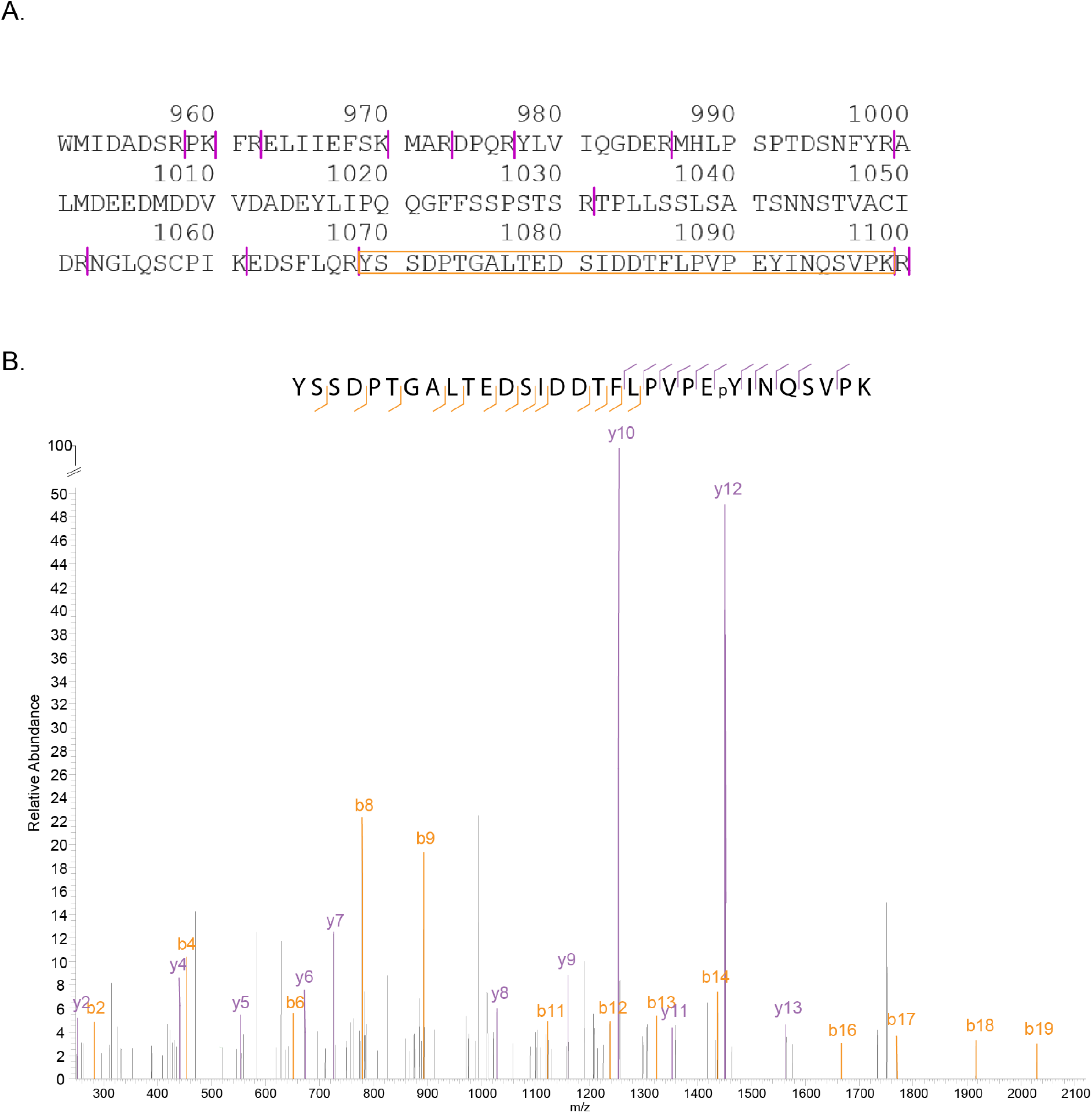
Mass spectrum of the Y1068 EGFR phosphorylated activation site. (A) Partial amino acid sequence of the human EGFR receptor. Purple vertical lines indicate tryptic cleavage sites. The orange square indicates the tryptic peptide containing the Y1068 activation site. (B) Mass spectrum of the phosphorylated Y1068 EGFR activation site.

## Notes

### Competing Interest Statement

The authors have declared no competing interest.

